# Temporal fMRI Dynamics Map Dopamine Physiology

**DOI:** 10.1101/2025.03.24.645022

**Authors:** Ian C Ballard, Ioannis Pappas, Daniella J Furman, Anne S Berry, Blaise deB Frederick, Robert L White, Andrew S Kayser, William J Jagust, Mark D’Esposito

## Abstract

Spatial variations in dopamine function are linked to cognition and substance use disorders but are challenging to characterize with current methods. Because dopamine influences blood vessel dilation, we hypothesized that hemodynamic latency, which reflects BOLD signal timing, could serve as an indirect marker of dopamine physiology. Across four datasets, we found a topography of hemodynamic latencies that precisely distinguished the nucleus accumbens, a dopaminergic region implicated in motivation and substance abuse, from other striatal regions. Using PET, genetics, and pharmacology, we found that hemodynamic latencies are robustly related to dopamine function and dopamine-linked behavior. In individuals with cocaine use disorder, we observed a spatial gradient of altered hemodynamic latencies in the striatum. This pattern independently predicted nicotine use, revealing a conserved physiological profile associated with addictive substance use. Hemodynamic latencies map regional, individual, and pathological differences linked to dopamine, opening new avenues for indirectly assessing the role of dopamine in healthy cognition and disease.

## INTRODUCTION

The neurotransmitter dopamine is involved in an array of cognitive and motor processes, including associative learning^1^, working memory^2^, motivation^3^, and decision-making^4^. Dysregulation of the dopamine system is implicated in many psychiatric^5–7^ and neurological disorders^8,9^. Consequently, tools for assessing dopamine physiology in humans can advance our understanding of healthy cognition and disease processes. However, techniques such as positron emission tomography (PET) have limited spatial resolution, which makes it challenging to profile spatial variations in dopamine physiology within individual brain regions. We present a new approach for indirectly probing spatial patterns of dopamine physiology with fMRI and use this method to reveal a gradient of altered striatal physiology in cocaine use disorder.

Research in animal models has shown regional differences in dopamine physiology, including release, reuptake, and receptor function, in the nucleus accumbens (NAcc) relative to other striatal nuclei^10,11^. This unique physiology of NAcc dopamine is thought to underlie its role in learning, motivation, and the initial reinforcing properties of addictive substances^11^. Approaches for defining dopaminergic regions based on neuroanatomical markers or structural and functional connectivity do not measure such regional variations in dopamine physiology^12–15^. New imaging techniques are needed to indirectly assess this spatial variation in striatal dopamine physiology in humans.

Although fMRI cannot directly measure dopamine or its physiology, dopamine release influences the blood-oxygen-level-dependent (BOLD) response^16^. There is a relationship between dopamine and the *magnitude* of the BOLD response to reward in the striatum or dopaminergic midbrain. These responses are correlated with reward prediction error^17,18^, a signal conveyed by dopamine neurons^1^, and co-vary with individual differences in dopamine physiology^19,20^. However, striatal prediction error BOLD responses do not correspond strongly to dopamine measured with voltammetry^21^. Additionally, the spatial location of BOLD responses is highly dependent on task variables (e.g., primary versus secondary rewards^17,18,22^), which makes it impossible to disentangle spatial differences in dopamine physiology from spatial differences in grey matter activation. For these reasons, event-related fMRI is poorly suited to assess spatial variations in dopamine physiology.

Emerging evidence supports the hypothesis that the *timing* of the BOLD signal is influenced by local dopamine neurophysiology. Dopamine varicosities are close to arterial walls, and dopamine axons have been observed to wrap around arterial microvessels^23^. Dopamine administration causes vasoconstriction^24^, and dopaminergic drugs influence arterial dilation^23–25^ and blood flow^26–28^. Because vasoconstriction is directly related to the velocity of blood flow^29^, higher extracellular dopamine should reduce the speed with which arterial blood arrives in tissue. The latency of the hemodynamic signal is sensitive to blood arrival times and can be measured in the absence of a task across striatal gray matter^30^. Given that extracellular dopamine transients persist longer in the NAcc compared with striatal regions^31^, we predicted increased hemodynamic latencies in the NAcc that correlate with individual and pathological increases in dopamine function.

In four fMRI datasets, we found a spatial gradient of hemodynamic latencies in the striatum that trace the anatomical boundary of the NAcc. Striatal hemodynamic latencies were correlated with multiple measures of dopamine physiology, causally altered by dopamine pharmacological manipulation, and linked to behavioral perseveration. Additionally, we found a spatial gradient of altered striatal hemodynamic latencies in cocaine use disorder. Our results show that hemodynamic latency can be used to indirectly probe dopamine physiology sensitive to regional, individual, and substance-abuse-related differences in dopaminergic function.

## RESULTS

We first asked whether hemodynamic latency was sensitive to regional differences in dopamine physiology in the striatum. We focused on the anatomical boundary separating the NAcc from other striatal regions, including the caudate and putamen. Although there are no anatomical markers of this boundary^32^ (Figure 1A), there are significant differences in dopamine physiology in the NAcc relative to other striatal regions, including differential baseline dopamine release dynamics^33,34^ and D2-receptor properties^35^, that result in slower fluctuations in extracellular dopamine^31^.

**Figure 1:**
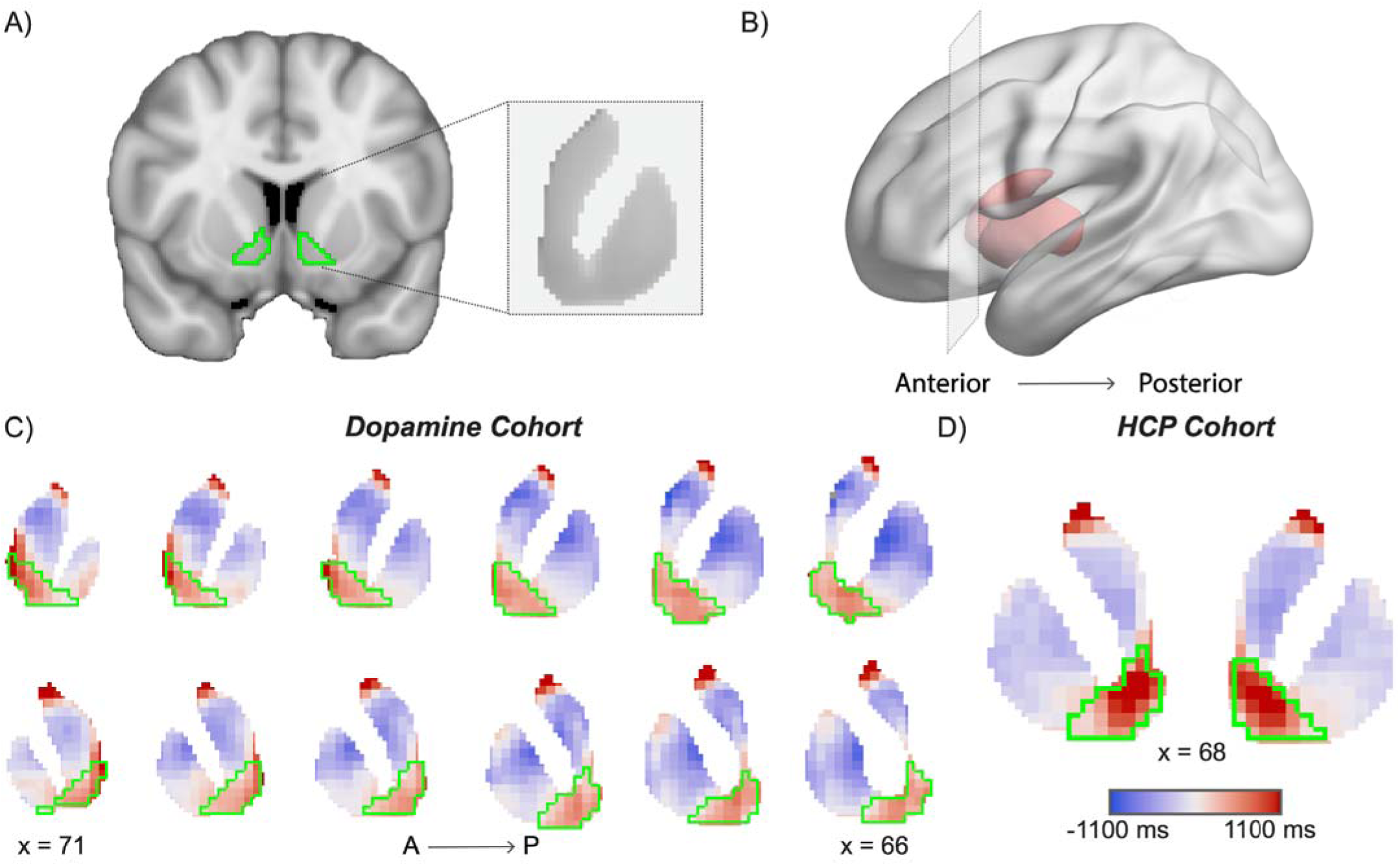
Hemodynamic latencies trace the anatomical boundary of the nucleus accumbens (NAcc). **A)** The anatomical outline of the NAcc (Harvard-Oxford atlas) is plotted in green. In the inset, note the absence of gross structural markers of the boundary between the NAcc and other striatal regions. **B)** Depiction of coronal sections through striatum (red) ranging from anterior to posterior. **C)** Hemodynamic latencies from the cohort with MRI and PET data are shown on coronal slices of the striatum. Higher latencies (red) indicate that the hemodynamic signal in that voxel lags the whole-brain signal. The light green outline of the anatomical NAcc boundary corresponds to a transition from positive to negative hemodynamic latencies. **D)** Hemodynamic latencies from the human connectome project (HCP) cohort replicate the close correspondence between hemodynamic latency and the NAcc anatomical boundary. *Coordinates are in MNI space*.

Healthy subjects (*n* = 73) underwent fMRI imaging following ingestion of a placebo, bromocriptine (a D2 agonist), or tolcapone (a brain-penetrant catechol-O-methyltransferase [COMT] inhibitor) in a within-subject, double-blind study performed across three sessions in randomized order. Using data from the placebo session, we created a hemodynamic latency map by comparing the timing of the BOLD signal in each voxel to the average signal from the whole brain using a temporal cross-correlation approach following established methods (See *Methods*)^36,37^. Positive latency in a voxel indicates that its signal lags the whole brain signal, whereas a negative latency indicates that its signal precedes the whole brain signal. This analysis revealed a striking correspondence between the hemodynamic latency map and the anatomical boundaries of the NAcc derived from the publicly available Harvard-Oxford anatomical atlas (Figure 1A-C).

Quantitatively, we found that hemodynamic latencies were substantially increased in the NAcc relative to the caudate, *Z =* 8.4, p < .001, Δlatency = 545 ms, 95% *CI* = [418, 672], and putamen, *Z =* 14.6, p < .001, Δlatency = 948 ms, 95% *CI* = [821, 1075]. Hemodynamic latencies in the striatum decreased linearly as the distance from the NAcc boundary in the striatum increased, *Z =* -11.8, p < .001, Δlatency/3mm voxel = -285 ms, 95% *CI* = [-238, -332] (Figure 2A). The sharp change in hemodynamic latency at the NAcc boundary, with latency decreasing by 95 ms/mm past the boundary, suggests a significant influence of NAcc physiology on hemodynamic latency. To ensure this pattern was not idiosyncratic to our dataset, we repeated the latency analysis in a large Human Connectome Project (HCP) sample and found a remarkably similar spatial pattern (Figure 1D).

**Figure 2:**
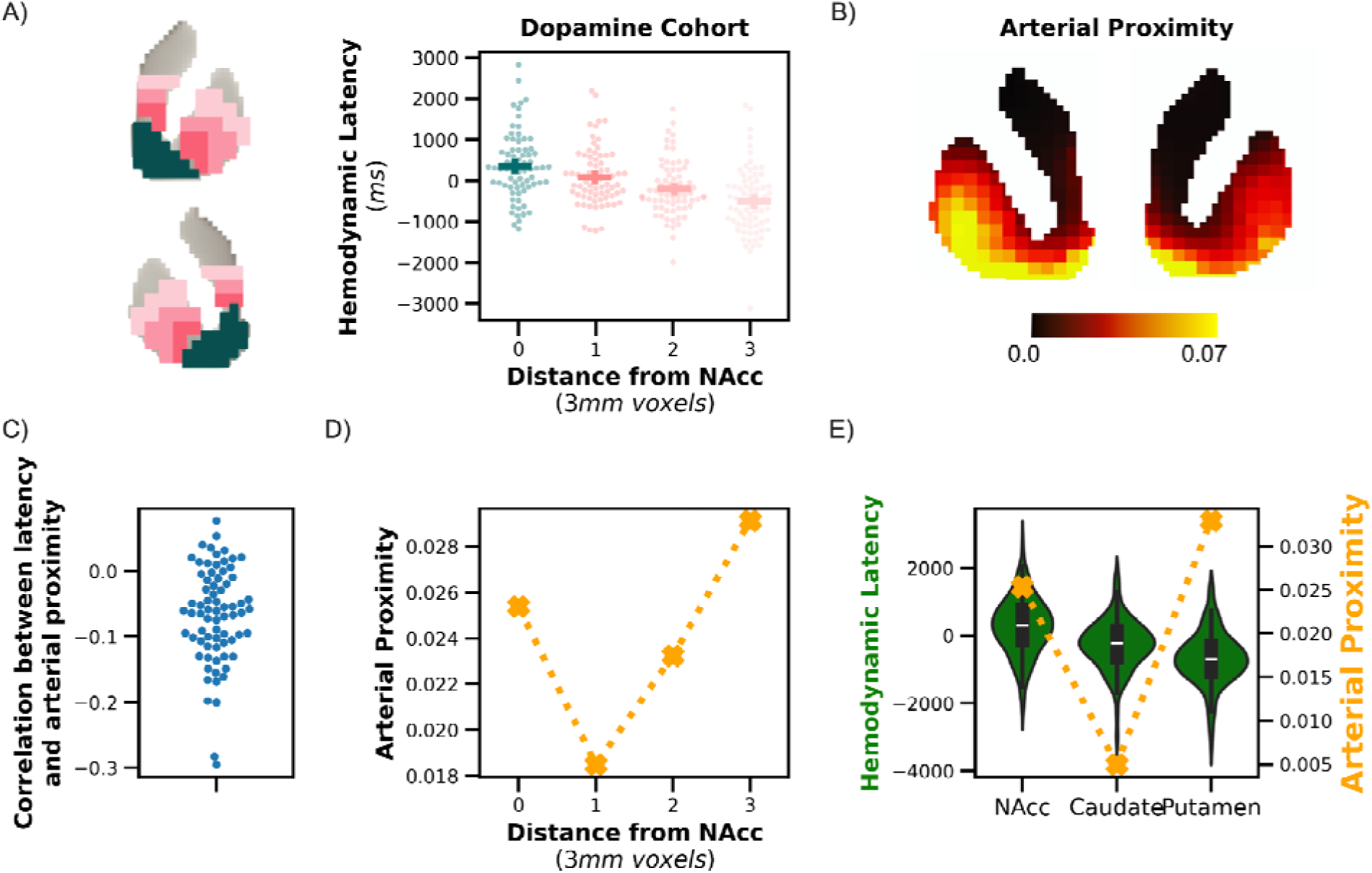
The spatial correspondence between hemodynamic latency and the NAcc boundary is not driven by proximity to major arteries. **A)** Hemodynamic latency in the striatum decreases linearly as distance from the NAcc increases. **B)** Spatial map of estimated proximity to major arteries in the striatum. **C)** Hemodynamic latency is negatively correlated with proximity to major arteries across whole-brain gray matter. **D)** Arterial proximity shows a distinct spatial profile from hemodynamic latency (compared with A). **E)** Differences in hemodynamic latency across striatal subregions cannot be explained by arterial proximity. Medians and quartiles of hemodynamic latencies are plotted as box plots within violin plots. *Error bars depict the standard error of the mean (SEM)*

Because hemodynamic latencies in major brain arteries are reduced relative to those in grey matter^38^ and spatial blurring of the BOLD signal could cause the signal from major arteries to blend into nearby grey matter, we tested whether differential proximity to major arteries could explain the spatial profile of hemodynamic latencies we observed in the striatum. However, arterial proximity showed a markedly different spatial distribution in the striatum compared to hemodynamic latency (Figure 2A, B). Across the brain, we found a small but significant correlation between the proximity of gray matter to major arteries and hemodynamic latency, *Z =* -7.7, p < .001, average *r* = -.067, 95% *CI* = [-.085, -.05] (Figure 2C). In a model of hemodynamic latency that included distance from the NAcc boundary and arterial proximity, we did not find an influence of arterial proximity, *p* > .2, whereas the influence of distance from the NAcc boundary remained large, *Z =* -10.3, p < .001, Δlatency/3mm voxel = -280 ms, 95% *CI* = [-227, -333]. Moreover, the regional pattern of hemodynamic latencies does not match the regional pattern of arterial proximity (Figure 2E).

We also examined whether proximity to large veins influenced hemodynamic latency estimates. We downloaded publicly available resting state fMRI from the Midnight Scan Club dataset^39^, which, unlike our other datasets, includes individual venograms and angiograms. We constructed a noise model that accounts for the spatial blurring of hemodynamic latencies from white matter, ventricles, and large veins and arteries into grey matter^38,40^ (Figure S1A). After scrubbing the data of these influences, hemodynamic latencies again demarcated the NAcc boundary, *Z =* -7.2, p < .001, Δlatency/3mm voxel = -223 ms, 95% *CI* = [-284, -162] (Figure S1B,C), indicating that the demarcation of the NAcc by hemodynamic latencies is not due to effects of spatial proximity to major sources of physiological noise. In additional exploratory analyses, we found evidence that increased hemodynamic latencies observed in the dorsal caudate, but not the NAcc, are driven by proximity to deep cerebral white matter (*Supplemental Text*, *Figure S2*).

Finally, we assessed whether the arterial supply of striatal gray matter could influence hemodynamic latencies. The NAcc is supplied by a branch of the anterior cerebral artery (ACA), whereas the dorsal striatum is supplied by branches of the middle cerebral artery (MCA)^41^. We measured the latency of all grey matter voxels within the vascular territories of the ACA and MCA, derived from the John Hopkins University (JHU) atlas, in the placebo session of the dopamine dataset^42^. We did not find a difference in latency between these vascular territories, *p* = .12. Thus, the vascular supply of the striatum is not likely to account for the observed pattern of hemodynamic latencies.

### Dopaminergic relationships to hemodynamic latency

We next examined whether hemodynamic latencies were sensitive to physiological markers of dopamine (Figure 3A). A subset of subjects (*n* = 44) underwent three positron emission tomography (PET) scans to assess distinct aspects of dopamine physiology: dopamine synthesis capacity (*K_i_*) with [^18^F]fluoro-l-m-tyrosine (FMT) and D2/3 receptor availability with [^11^C]raclopride (raclopride has affinity to both D2 and D3 receptors), and dopamine release using [^11^C]raclopride displacement with methylphenidate. We found that subjects with higher presynaptic dopamine synthesis capacity in the NAcc exhibited reduced hemodynamic latencies in the NAcc on placebo, *Z =* -3.0, p = .003, Δlatency/fmt *K_i_* =-145,000, 95% *CI* = [-241,000, -50,000] (Figure 3B). This relationship was not altered by either dopaminergic drug, *p* > .2.

**Figure 3:**
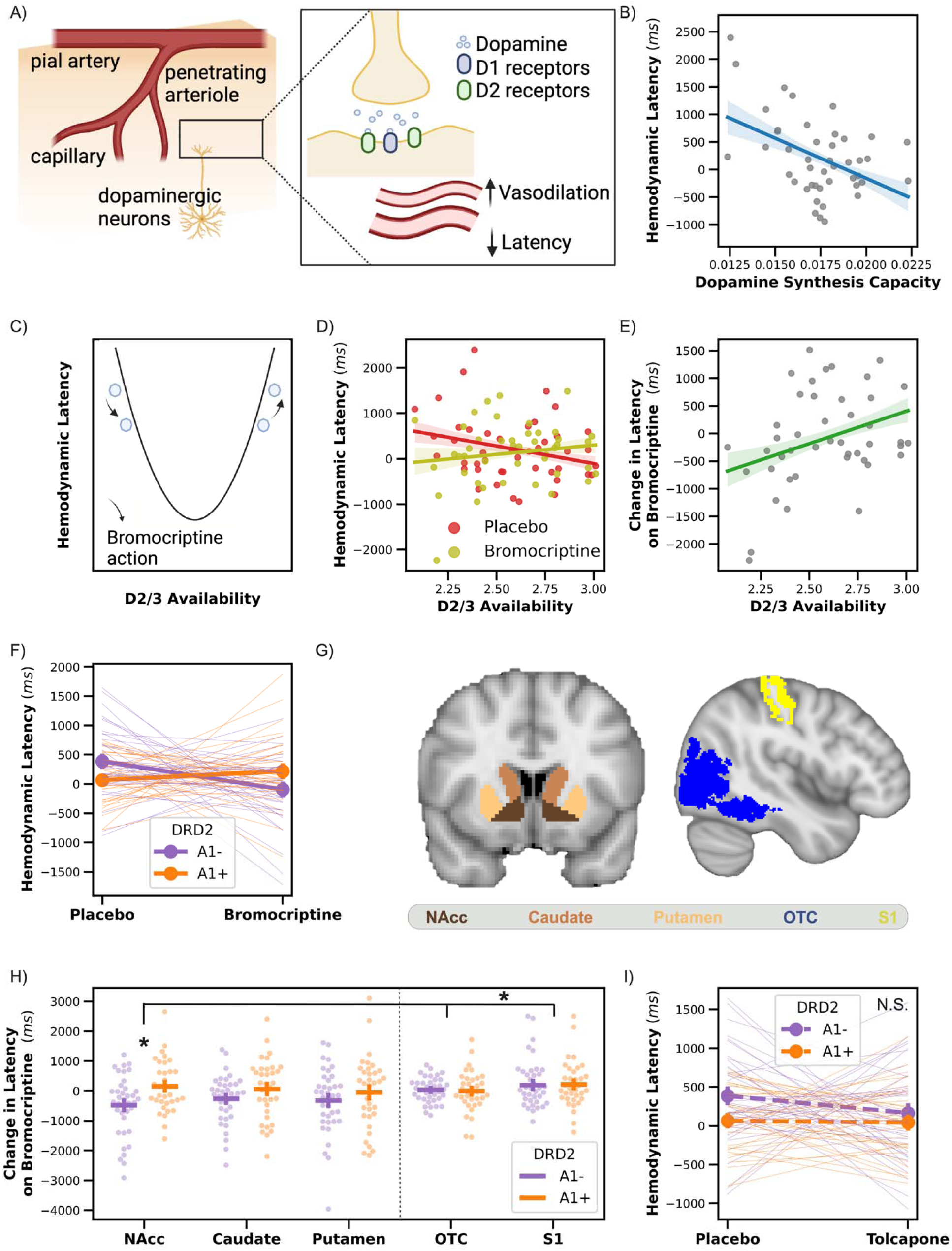
Hemodynamic latencies are influenced by dopamine physiology. **A)** Dopamine release influences vasodilation, which should be reflected in altered hemodynamic latency. **B)** Subjects with higher dopamine synthesis capacity in the NAcc have reduced hemodynamic latencies on placebo. **C)** The U-shaped relationship between D2 activation and hemodynamic latency suggests that administration of bromocriptine, a D2 agonist, could have opposing influences on hemodynamic latency depending on baseline D2/3 availability. **D)** The relationship between D2/3 availability and hemodynamic latencies is altered with bromocriptine administration. **E)** Bromocriptine reduces hemodynamic latency more strongly for subjects with lower D2/3 availability in the NAcc, consistent with the U-shaped model. **F)** Bromocriptine reduces hemodynamic latency more for individuals with the DRD2 A1-allele. **G)** Anatomical definitions of regions of interest, including striatal subregions enriched in dopamine and non-dopaminergic cortical regions. **H)** The interaction between bromocriptine and DRD2 genotype is stronger in the NAcc than in non-dopaminergic cortical regions. The dashed line separates striatal subregions (left) from non-dopaminergic cortical regions (right). **I)** Administration of tolcapone, a COMT inhibitor that does not directly influence D2 receptors, does not interact with the DRD2 allele to influence hemodynamic latency. *OTC: occipitotemporal cortex; S1: somatosensory cortex*. * marks a significant interaction; *NS* indicates not significant. *Error bars indicate SEM*.

Our finding of increased hemodynamic latencies in the NAcc relative to other striatal subregions could reflect regional differences in dopamine physiology. To test this hypothesis, we asked whether the relationship between dopamine synthesis capacity and hemodynamic latency differed across striatal subregions. The inverse relationship between dopamine synthesis capacity and hemodynamic latency was more pronounced in the NAcc than in the caudate, *Z =* 3.2, p = .001, Δlatency/fmt *K_i_* = 216,000, 95% *CI* = [83,000, 350,000], and putamen, *Z =* 2.7, p = .007, Δlatency/fmt *K_i_* = 176,000, 95% *CI* = [49,000, 303,000]. Only in the NAcc was there a significant negative relationship between dopamine synthesis capacity and latency, and this relationship was numerically reversed in sign in the caudate, *Z =* 1.7, p = .089, Δlatency/fmt *K_i_* = 104,000, 95% *CI* = [-16,000, 223,000]. Therefore, hemodynamic latencies show distinct relationships with dopamine synthesis capacity in striatal regions with distinct dopamine physiology and timescales of dopamine function^31^.

In slice preparations, pharmacological activation of D2 receptors drives vasodilation in a dose-dependent, U-shaped manner^24^. Therefore, we predicted that pharmacological activation of D2 receptors with bromocriptine would influence hemodynamic latencies differently depending on baseline D2/3 receptor availability (Figure 3C). Consistent with the U-shape model, bromocriptine reduced hemodynamic latencies in the NAcc more strongly for lower D2/3 availability subjects, bromocriptine X D2/3 availability interaction: *Z =* 2.2, p = .027, *B* = 1176, 95% *CI* = [132, 2220] (Figure 3D, E), with no main effects of bromocriptine, *p* > .2, or D2/3 receptor availability, *p* = .085. The interaction between bromocriptine and D2/3 availability was not significantly different between the NAcc and caudate, region X bromocriptine X D2/3 availability *p* = .063, or putamen, region X bromocriptine X D2/3 availability *p* = .068. Thus, both high and low levels of D2 receptor activity increase hemodynamic latencies across the striatum. Finally, we did not find a relationship between dopamine release and hemodynamic latency, *p* > .2.

### Dopamine genetic relationships to hemodynamic latency

The Taq1A polymorphism of the DRD2 gene is associated with altered D2 receptor function and increased propensity for addictive behaviors^43^. We examined whether the Taq1A polymorphism showed a U-shaped relationship with hemodynamic latencies (Notably, there was no difference in D2/3 receptor availability between Taq1A groups in our sample, *p* > .2). Consistent with the U-shaped model, bromocriptine reduced latencies more strongly for A1-carriers than A1+ carriers in the NAcc, bromocriptine X DRD2 interaction *Z =* 2.9, *p* = .004, Δlatency = 316 ms, 95% *CI* = [103, 530] (Figure 3F), and this interaction was not different between striatal ROIs, bromocriptine X DRD2 region interaction *p*s > .2. Neither bromocriptine, *p* = .13, nor Taq1A genotype, *p* = .068, influenced hemodynamic latencies.

Because D2 receptors could influence overall brain vascular function, we assessed whether the influence of bromocriptine on hemodynamic latency varied by DRD2 genotype in regions without substantial dopamine innervation: the occipitotemporal cortex (OTC) and the somatosensory cortex (S1)^44^. Critically, there was a triple interaction of region, drug, and genotype, such that the interaction between DRD2 and bromocriptine was larger in the NAcc than OTC, *Z =* -2.2, *p* = .025, β = -337, 95% *CI* = [-632, -42], and S1, *Z =* -2.0, p = .043, β = -305, 95% *CI* = [-600, -10] (Figure 3H), indicating that D2 receptor properties are selectively related to striatal hemodynamic latencies.

It is possible that general stimulation of the dopamine system, rather than D2 agonism specifically, alters hemodynamic latencies. To test this, we administered tolcapone to the same subjects as a positive control. Unlike bromocriptine, tolcapone enhances cerebral extracellular dopamine by inhibiting dopamine clearance but is thought to have a weaker impact on striatal dopamine than other regions^45–47^.

Tolcapone did not influence hemodynamic latencies nor interact with D2/3 availability or DRD2 Taq1A polymorphism, *ps* > .2 (Figure 3I), confirming that agonism of D2 receptors has a specific, baseline-dependent influence on NAcc hemodynamic latency. In contrast, we found that tolcapone influenced hemodynamic latencies dependent on the COMT allele (Figure S3).

### Hemodynamic latency and perseverative behavior

Dysfunction in the dopamine system is associated with failures to update ongoing behavior appropriately^48^, and such perseveration is increased in drug abuse disorders^49^. In our study, subjects completed a rule-updating task requiring them to switch between number judgment rules (“odd/even” or “greater/less than 5”) based on intermittent switch cues (Figure 4A)^50^. Our previous research found that bromocriptine administration influenced perseverative errors—failures to switch rules after switch cues—in a manner that depended on the DRD2 genotype, implicating striatal dopamine in these errors^51^.

**Figure 4:**
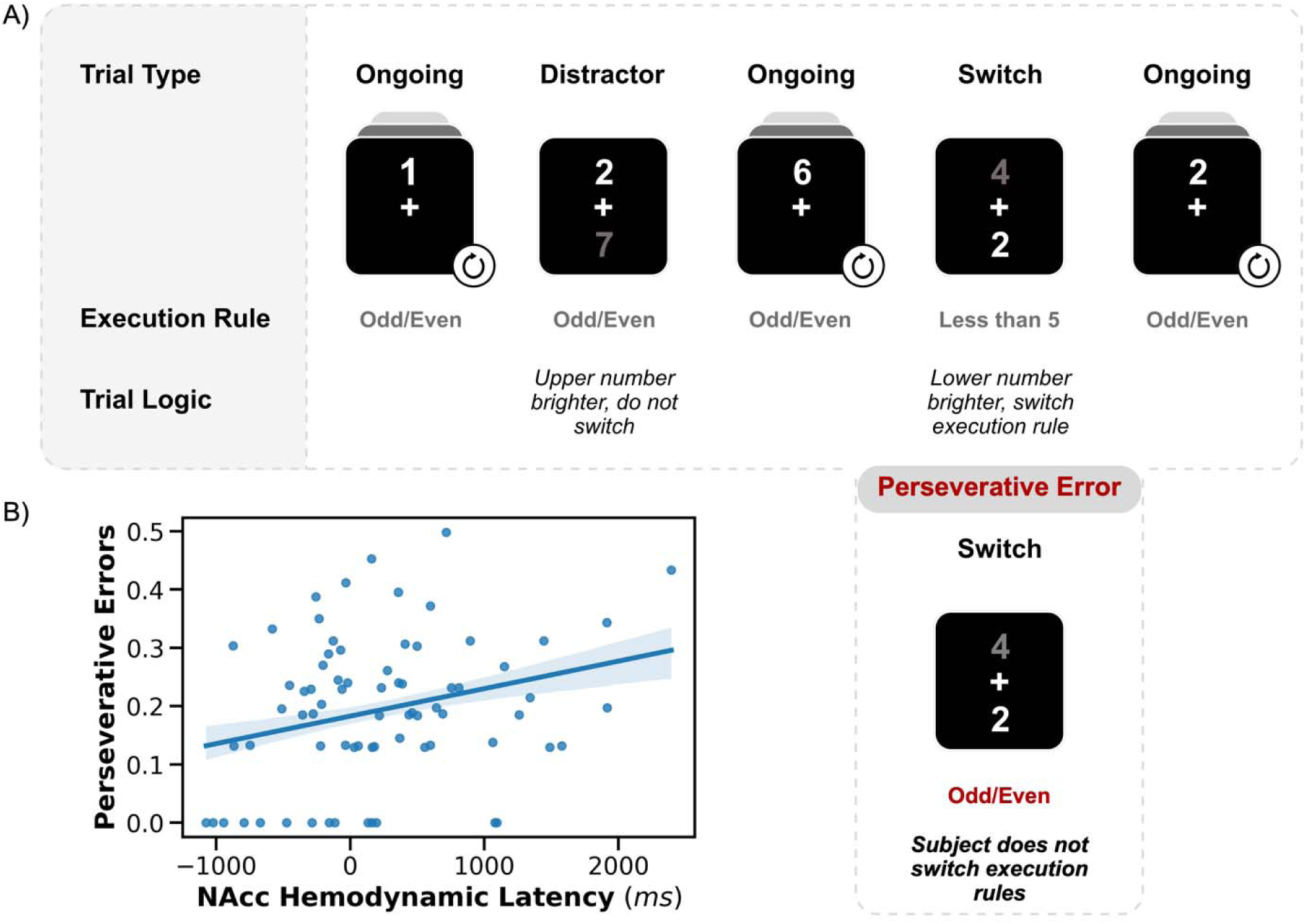
Cognitive task. **A)** In ongoing trials, subjects applied the dominant task rule (pictured: “odd/even”). When two numbers appeared above and below fixation, subjects responded differently depending on the location of the brighter number. If the upper digit was brighter (“distractor trials”), subjects continued to apply the dominant rule to the upper digit. If the lower digit was brighter (“switch trials”), subjects applied the alternative rule to the lower digit using different response keys (pictured: “less than 5”). *Perseverative errors* occur when the subject fails to switch rules on a switch trial. *Ambiguous trials*, where the two numbers are similarly bright, are not shown. **B)** NAcc hemodynamic latencies correlate with perseverative behavior. Error rates are arcsine-square-root transformed (*mean perseverative error rate* = 4.6%). *Error bars indicate S.E.M*.

Based on our findings that increased hemodynamic latency in the NAcc reflects reduced dopamine function, we predicted that higher latency would be associated with more frequent perseverative errors. Hemodynamic latency in the nucleus accumbens was positively associated with the perseverative error rate, *Z =* 3.0, FDR-corrected *p* = .016, *β* = 1950, 95% *CI* = [677, 3224], mean perseverative error rate = 4.6%, *n* = 73 (Figure 4B), even when controlling for the previously reported bromocriptine by DRD2 interaction, *Z* = 2.8, *p* = .005, *β* = 1827, 95% *CI* = [556, 3100]. This finding suggests that increased nucleus accumbens hemodynamic latencies may be a marker of dopamine-related behavioral deficits.

### Hemodynamic latency in cocaine use disorder

Our finding of increased hemodynamic latency in the NAcc is notable because altered dopamine physiology in the NAcc is related to the initial reinforcing properties of dopaminergic drugs^10,52,53,54^. Cocaine use disorder is associated with physiological adaptations, including internalization of D2 receptors, reduction of D2/3 availability, and elevated dopamine transporter function^55–58^ Given our finding that increased hemodynamic latency in the NAcc reflect reduced dopamine function, we hypothesized that hemodynamic latencies in the NAcc would be elevated in individuals with cocaine use disorder. We calculated hemodynamic latency maps from a publicly available dataset of individuals with cocaine use disorder and a control group matched for age, sex, handedness, and education^59,60^.

We found that hemodynamic latencies in the NAcc were substantially increased in the cocaine use disorder group relative to the control subjects, *Z =* 3.5, p = .001, Δlatency = 789 ms, 95% *CI* = [341, 1237] (Figure 5A). Additionally, we replicated the finding that hemodynamic latency rapidly decreases beyond the NAcc boundary, *Z =* -6.8, p < .001, Δlatency/3mm voxel = -201 ms, 95% *CI* = [-259, -143] (Figure S4).

**Figure 5:**
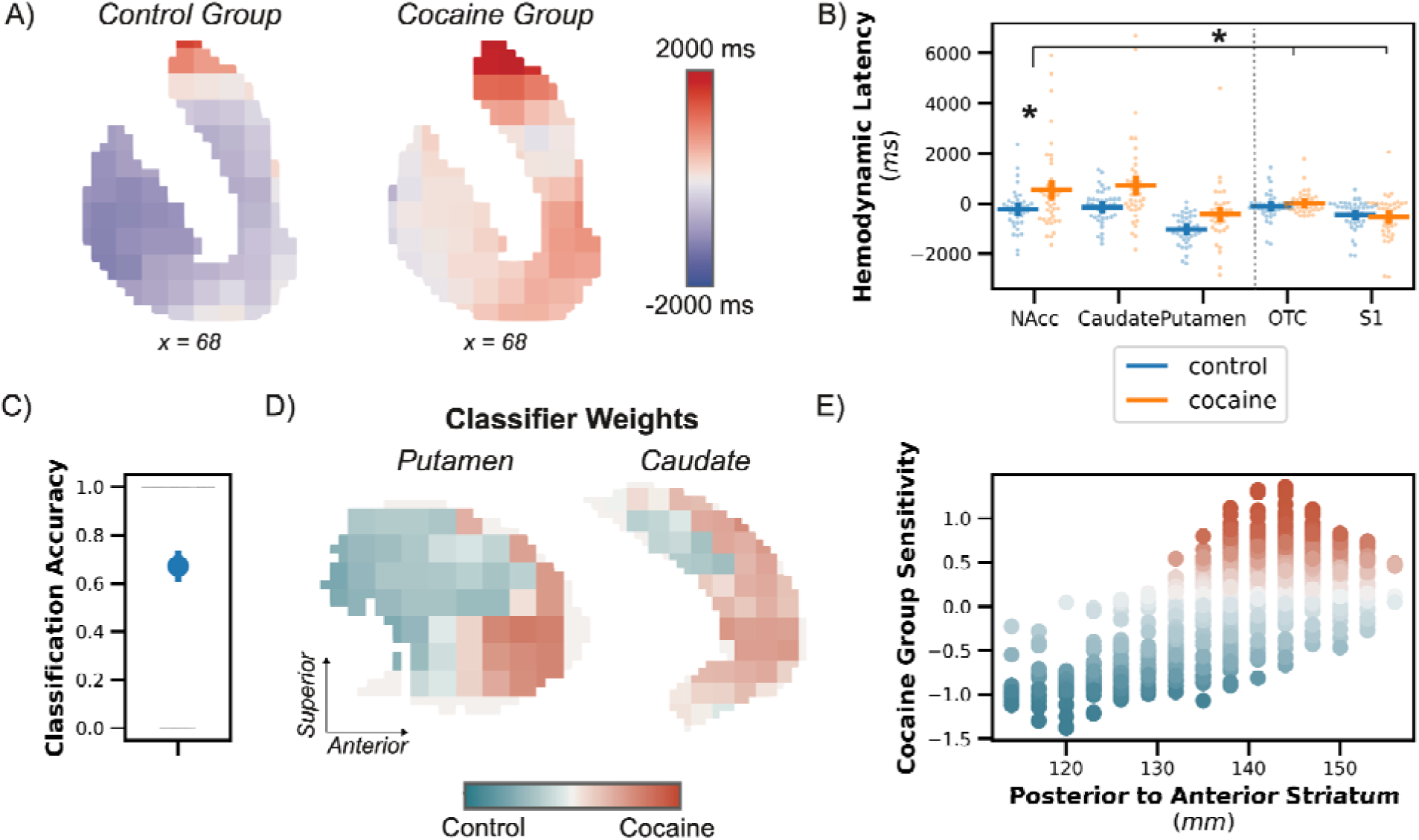
Hemodynamic latencies in cocaine use disorder. **A)** Hemodynamic latencies are increased in the striatum for subjects with cocaine use disorder relative to controls. Note that the scale obscures the significant difference in latency between nucleus accumbens and dorsal striatum in both groups; see *Figure S4*. **B)** Hemodynamic latencies increase more strongly in the NAcc in subjects with cocaine use disorder than in non-dopaminergic cortical regions. The dashed line separates striatal subregions (left) from non-dopaminergic cortical regions (right). **C)** Classification accuracy for a classifier trained to predict cocaine use or control group status from the spatial patterns of hemodynamic latencies in the striatum. **D)** Spatial pattern of importance weights from the classifier plotted on the putamen and caudate. **E)** Increased hemodynamic latency in anterior portions of the striatum is more predictive of cocaine use. * marks a significant interaction. *Error bars depict the SEM*.

Because cocaine use disorder could influence overall brain vascular function^61^, we assessed whether the difference in latency between groups was similar in non-dopaminergic cortical regions. We found that the difference in hemodynamic latencies between the groups was larger in the NAcc than either OTC, *Z =* -2.4, p = .015, *B* = -652 ms, 95% *CI* = [-1177, -126], or S1, *Z =* -3.3, p = .001, *B* = -871 ms, 95% *CI* = [-1396, -346] (Figure 5B). Thus, cocaine use disorder is associated with an increase in hemodynamic latency in the striatum without a concomitant change in cortical regions lower in dopamine.

We did not observe regional differences between cocaine and control groups across striatal subregions, *p* > .2. Caudal striatal regions linked to habit formation become involved later in the course of cocaine use disorder and have less pronounced physiological adaptation^62^. Therefore, we hypothesized that there would be an anterior to posterior gradient of altered hemodynamic latencies associated with cocaine use disorder. We built a classifier to predict cocaine use disorder based on striatal hemodynamic latencies. Importantly, we removed the mean hemodynamic latency across striatal voxels within each subject, ensuring that our classifier predicted cocaine use disorder solely based on the spatial pattern of hemodynamic latencies. Our classifier predicted cocaine use disorder with 67.1% cross-validated accuracy, *permuted p* = .016 (Figure 5C).

To visualize the topography of altered hemodynamic latencies in cocaine use disorder, we examined the spatial pattern of importance weights from our classifier. Larger importance weights indicate that increased hemodynamic latency is predictive of cocaine use disorder. Consistent with our prediction, we observed a gradient across the striatum whereby hemodynamic latencies in more anterior voxels were more strongly linked to the probability of cocaine use disorder, *B* = .43, *permuted p* = .027 (Figure 5D, E). We did not find evidence of a medial-to-lateral or dorsal-to-ventral spatial gradient of importance weights, *p* > .2.

Because nicotine also increases striatal dopamine release^34,52^, we assessed whether the pattern of hemodynamic latencies that predicts cocaine use disorder would generalize to tobacco use. Because few control subjects used tobacco, we restricted our analysis to tobacco users in the cocaine use disorder group. Subjects who were more likely to have cocaine use disorder based according to the classifier log probabilities also had a higher level of daily tobacco use (Modal tobacco use: 1-5 cigarettes/day, range: 1 to >21), *spearman r*(33) *=* .37, *p* = .025. There was no relationship between the level of daily tobacco use and weekly cocaine use, *spearman r*(33) = .02, *p* > .2, nor was there a relationship between classifier probabilities and age, *spearman r*(33) = -.12, *p* > .2. Although the sample size for this analysis indicates caution in interpretation, the generalization of the cocaine use disorder classifier to nicotine use suggests that increased hemodynamic latencies in the anterior striatum reflect addiction-related differences in dopamine physiology.

## DISCUSSION

Using dense phenotyping of individual neurophysiology, we found convergent and causal evidence that hemodynamic latencies in the striatum reflect dopamine neurophysiology. Specifically, higher hemodynamic latencies in the NAcc were associated with lower dopamine synthesis capacity, more frequent perseverative errors, and distinguished cocaine use disorder subjects from control subjects. Perseverative errors and cocaine use disorder are both associated with reduced dopamine function^48,58^, and perseveration is elevated in cocaine use disorder ^49^ . These findings suggest that elevated hemodynamic latencies in the NAcc are an indirect marker of reduced dopaminergic function.

D2 receptors are linked to learning, decision-making, and behavioral control^63–65^. In slice preparations, both D2 agonists influence vasodilation in a U-shaped manner: moderate doses cause vasodilation, whereas low or high doses cause vasoconstriction^24^. These results are mirrored by our finding that bromocriptine, a D2 agonist, causally influenced striatal hemodynamic latencies differently depending on D2/3 availability, with increased latency associated with both high and low D2 activity. Similarly, the impact of bromocriptine on striatal hemodynamic latencies depended on the Taq1A allele of the DRD2 gene, which encodes the D2 receptor. Hemodynamic latencies are informative about D2 receptor function and can, when combined with pharmacological manipulations, contribute to an improved understanding of D2-linked behaviors.

Hemodynamic latencies exhibited a spatial topography sensitive to differences in striatal dopamine physiology. Hemodynamic latencies were substantially increased in the nucleus accumbens and precisely demarcated its anatomical boundary. Moreover, the relationship between dopamine synthesis capacity and latency differed between striatal regions, potentially reflecting different timescales of extracellular dopamine^31^.

We also found that cocaine use disorder and nicotine use were associated with a shared anterior-to-posterior gradient of increased hemodynamic latencies in the striatum. These results highlight that hemodynamic latencies reveal spatially specific information linked to local dopamine neurophysiology.

PET is the gold standard in assessing human dopamine physiology because it measures specific markers, including presynaptic dopamine synthesis capacity, D2/3 receptor availability, and others^66^. However, PET has limited spatiotemporal resolution^67,68^ due to imaging constraints. Thus, we are unaware of any PET ligand that sharply distinguishes the NAcc from the dorsal striatum. More importantly, hemodynamic latencies revealed distinct information from our PET assays: Latencies were influenced by multiple aspects of dopamine physiology, and the relationship between dopamine synthesis capacity and latency differed between striatal subregions. Thus, although hemodynamic latencies are indirect assays and cannot match the mechanistic specificity afforded by PET, they could provide complementary information about the combined impact of multiple dopamine synthesis, release, and reuptake mechanisms.

Striatal dopamine levels fluctuate over multiple timescales, from seconds to hours^69–71^, and these fluctuations are linked to different timescales of learning and reward expectation^31^. Future work is needed to determine whether fluctuations in hemodynamic latencies within an individual are linked to dopamine signals like reward prediction error. Additionally, whether altered hemodynamic latencies in cocaine use disorder normalize in remission is an open question. Hemodynamic latencies are a novel, indirect proxy of dopamine with the potential to reveal dopamine substrates of cognitive function in healthy individuals and disorders such as substance abuse and Parkinson’s disease.

## METHODS

### Hemodynamic latency

It has been shown that low-frequency oscillations (∼0.01-0.1Hz) in the fMRI signal are associated with blood circulation^72^. The delay of these low-frequency fMRI signals correlates with the time it takes for blood to propagate to different brain regions ^30^. To extract these latencies, we used the RIPTiDe processing stream (https://github.com/bbfrederick/rapidtide). First, the fMRI data is bandpass filtered at 0.009-0.15Hz^36^. Second, for each voxel, hemodynamic latencies are calculated as the delay yielding the highest cross-correlation between the delayed time series in that voxel and the global time series (in our case, the average signal within the brain mask)^73^.

The RIPTiDe algorithm iteratively refines the cross-correlation to avoid noisy latency estimations. After an initial cross-correlation, each voxel time series (excluding those with noisy latencies) is shifted based on their peak latency and averaged to create a new global time series. Latencies are calculated again via cross-correlation with the new global time series. We used three iterations to refine the global regressor, consistent with previous studies^37^. Finally, we extracted the median latency within our regions of interest. Striatal hemodynamic latency maps were resampled to 2 x 2 x 2 mm for visualization.

### Magnetic Resonance Angiography dataset

To assess the contribution of arterial signal to hemodynamic latency, we used a publicly available artery occurrence probability map generated based on magnetic resonance angiography images from 544 individuals^74^. We resampled the images to functional space and simulated the effect of spatial blurring of arterial signals into nearby gray matter by applying a Gaussian spatial filter to the probability maps with an 8 mm FWHM. The resulting smoothed probability images correspond to the probability of arterial signal given Gaussian spatial filtering. In our analyses, we refer to this quantity as *arterial proximity*.

### Dopamine dataset

The procedures for the dopamine dataset have been previously described in published work using these data^28,51,63^. All subjects (*n* = 77, 48 female sex, median age = 21, age range 18 - 30, standard deviation = 2.46 years) gave written, informed consent in accordance with the procedures approved by the Committee for the Protection of Human Subjects at the University of California, San Francisco, the University of California, Berkeley, and the Lawrence Berkeley National Laboratory and were compensated for their participation. One subject did not complete the placebo MRI session, three subjects had unusable MRI acquisitions^51^, leaving 73 subjects in the reported results. Of these, 44 subjects also underwent PET imaging.

#### Genotyping procedures

Saliva samples were obtained using Oragene collection kits with stabilizing liquid (DNA Genotek). Genotyping of COMT (rs4680) and Taq1A (rs1800497) SNP testing was performed at the UCSF Genomics Core, Vincent J. Coates Genomics Sequencing Laboratory, and Kashi Clinical Laboratories using polymerase chain reaction-based TaqMan technology (Applied Biosystems). Only individuals who were homozygous for either the Val or Met allele of the COMT polymorphism were invited to participate in the remainder of the study. Taq1A genotypes were binned according to the presence (“A1+”) or absence (“A1−”) of any copies of the A1+ minor allele. Subjects were selected based on compound COMT/Taq1A genotype, with roughly equal representation in each of the following groupings: Met/A1+ (n = 20), Val/A1+ (n = 17), Met/A1− (n = 21), Val/A1− (n = 19).

#### Cognitive task procedures

A detailed description of the task procedures and data cleaning is available in prior reports^51^, and the current study uses identical data treatment methods. While undergoing fMRI scanning, subjects responded quickly to digits between 1 and 9 (excluding 5) that appeared in shades of gray against a black background for 990 trials. Most trials were “ongoing task” trials (82%) where a single digit appeared above a central fixation cross. Subjects implemented a rule-based decision about the digit (either “odd/even” or “greater/less than 5” is the dominant rule for the block) and responded using the index finger of their left or right hand. Ongoing task trial types were presented in runs of three to six trials.

On the other 18% of trials, two digits were presented, one above and one below the fixation cross, and the relative brightness of the two digits cued the task rule. When the upper digit was brighter, subjects were instructed to ignore the lower digit and continue to apply the ongoing task rule to the upper digit (“distractor trials;” 6% of trials). When the lower digit was brighter, subjects were instructed to apply the alternate task rule to the lower digit (“switch trials;” 6% of trials). On switch trials, subjects were instructed to use their middle, rather than index, fingers to make responses, enabling us to distinguish between *perseverative errors* (continuing to use the ongoing rule and response with index fingers) and *execution errors* (incorrect responses using the correct rule). On the final third of trials (“ambiguous trials;” 6% of trials), the difference in brightness between the upper and lower digits was reduced. Digits were presented for 900 ms, but responses were recorded for the duration of the 2000 ms trial. There were no additional intertrial intervals.

We examined whether hemodynamic latencies in the nucleus accumbens were associated with six task variables: switch costs in accuracy, switch costs in response time, distractor costs in accuracy, distractor costs in response time, execution errors, and perseverative errors. As in prior reports, we applied arcsine-square-root transformations to error rates. We constructed separate random-intercept linear models of each task variable, with drug condition (treatment-coded, placebo baseline), nucleus accumbens hemodynamic latency, and their interaction as dependent variables. We applied FDR correction to the p-values of the six tests relating hemodynamic latency to task variables on placebo. Only perseverative errors were significantly related to hemodynamic latencies (all other *p*s > .2).

#### Pharmacology and fMRI procedures

Eligible subjects were scheduled for three pharmacological study sessions to be completed on different days. During the three sessions, subjects received a single oral dose of bromocriptine (1.25 mg), tolcapone (200 mg), or placebo in identical, compounded capsules. Drug doses were selected based on their ability to elicit changes in cognitive performance^75–77^.

In each drug session, subjects underwent fMRI imaging using a Siemens 3T Trio Tim scanner at UC Berkeley’s Brain Imaging Center. Anatomical data was obtained using T1-weighted magnetization prepared rapid gradient-echo (MPRAGE) scans [repetition time (TR) = 2300 ms; echo time (TE) = 2.98 ms; flip angle (FA) = 9°; bandwidth = 238 Hz/Pixel; matrix = 240 × 256; field-of-view (FOV) = 256 cm; sagittal plane; voxel size = 1mm^3; 160 slices. Functional MRI data was obtained during a task described elsewhere^78^. fMRI data were obtained during two 11-minute runs with a gradient-echo echo-planar imaging (EPI) sequence (TR = 2,000ms, TE = 24ms, flip angle = 65 degrees, field of view 224mm, 36 slices, AC-PC, voxel size = 3.0x3.0x3.5mm), 300 volumes. Preprocessing of the dopamine fMRI data has been described elsewhere^78^. Briefly, functional MRI data was slice-time and motion corrected and registered to anatomical data using 3dAllineate. Anatomical data was projected to MNI space using FNIRT, and the resulting transformations were used to map the coregistered functional data to MNI space.

We used residualized data obtained after modeling task variables for the latency analysis. Task-related regressions were constructed for each event using a spline basis set model. Distractor, switch, ambiguous, first, error, and miss onsets were modeled using CSPLIN in AFNI^79^. Motion traces, motion derivatives, and 0th-4th order trends were included in the model. Residuals were calculated using 3dDeconvolve in AFNI.

We extracted hemodynamic latencies from the NAcc, putamen, and caudate. To maximize correspondence to the PET data, we used the same hand-drawn ROIs used in PET data processing for the latency-PET correlation analyses^80^. For all other analyses, we aimed to maximize ease of reproducibility and relationship to existing literature and used public ROIs from the Harvard-Oxford atlas^81^. In addition, we defined two control ROIs from the primary sensory cortex, which receives a much lower density of dopamine projections. We defined a primary somatosensory (S1) ROI using a probabilistic atlas of the finger region of the postcentral gyrus (digitAtlas; all fingers combined)^82^. We also included a large control region to account for potential systemic effects on brain perfusion: the ventral occipitotemporal cortex (VTC). We defined this ROI by combining all category-selective areas of the VTC^83^. We used random-intercept mixed-effects models implemented in statsmodels to model hemodynamic latency as a function of genotype or PET measure, ROI, and ROI by genotype/PET interaction.

#### PET procedures

Fifty-two of the subjects completed three PET scans. The PET procedures and analysis techniques are described in prior reports using this dataset^80^. Briefly, subjects underwent a [^18^F]fluoro-l-m-tyrosine (FMT) PET scan to measure dopamine synthesis capacity, a [^11^C]raclopride PET scan to measure D2/3 receptor occupancy, and a [^11^C]raclopride PET scan one hour following a 30mg oral dose of methylphenidate (and then compared to the baseline scan) to measure dopamine release. All PET data were acquired using a Siemens Biograph Truepoint 6 PET/CT scanner. Data were reconstructed using an ordered subset expectation maximization algorithm with weighted attenuation, corrected for scatter, smoothed with a 4 mm full width at half maximum (FWHM) kernel, and motion corrected. Ventral striatal ROIs were hand-drawn on subjects’ T1-weighted images according to established procedures^84^. FMT data were analyzed using a reference region Patlak model^85^ to determine net tracer influx (K_i_) as the outcome variable of interest reflecting dopamine synthesis capacity. Raclopride analyses used a reference tissue reversible tracer mode^86^ to calculate non-displaceable binding potential (BP_ND_) as the outcome of interest reflecting receptor availability.

### HCP data

The rs-fMRI data from the Human Connectome Project (HCP) 1200 Subjects Release were used for this analysis^87^. In the HCP protocol, each subject underwent four ∼15-minute echo-planar imaging (EPI) resting state scans (3 Tesla Siemens Connectome Skyra, repetition time/echo time=720/33.1 milliseconds, multiband factor=8, 1200 volumes, 2 mm isotropic voxels, 72 slices covering the whole brain, FOV=208 x 180 mm): data were collected in two sessions on subsequent days; each session consisted of two acquisitions alternating between a left-right and a right-left phase encode direction. Our analysis included all subjects with complete data. Of the 1113 unique subjects in the release, 1009 met this criterion, yielding 4036 datasets.

The voxel-wise time course data that had gone through the volumetric “minimally preprocessed” analysis stream was used for this analysis^88^. This processing included the removal of spatial distortions, motion correction, alignment of the functional scan to the normalized anatomical scan, and normalization of time courses to the global mean^89,90^. This minimal preprocessing avoids removing signals needed for calculating hemodynamic latencies. Finally, an MNI152 brain mask was applied to restrict analyses to brain voxels.

### MSC dataset

The Midnight Scan Club (MSC) data were obtained from the OpenfMRI database ds000224. The data consists of structural and functional MRI data from 10 individuals (5 females, ages 24-34) scanned in ten sessions. Each session consisted of a resting-state scan and four task scans: a motor task, a semantic task, a coherence task, and an incidental encoding memory task. Detailed study procedures are reported in the original study^39^.

Functional and anatomical imaging data was processed using fMRIPrep^91–93^. Anatomical data was N4 bias-field corrected using N4BiasFieldCorrection. Then, normalization to the *ICBM 152 Nonlinear Asymmetrical template version 2009c* (Fonov et al. 2009, RRID: SCR_008796) was performed through nonlinear registration with antsRegistration (ANTs 2.2.0). BOLD data were minimally preprocessed to match the dopamine and HCP data. Specifically, data were motion-corrected and registered to the T1w image using a boundary-based cost function. BOLD data were resampled to MNI space using the BOLD-to-T1w and the T1w-to-MNI transformations. Motion parameters, their derivatives, and 0th-4th order trends were regressed from the functional data. Data was smoothed within a brain mask using FWHM of 4mm.

We constructed a noise model to remove the influence of hemodynamic latencies from non-grey matter on our estimates of striatal latencies. The MSC included venograms and angiograms for each subject. Venograms and angiograms were manually thresholded to isolate vessels and binarized, then resampled to match the resolution of the functional data. The noise model included the venogram, angiogram, white matter, deep white matter, and lateral ventricles. To account for the potential spatial blurring of signals from these tissue types into grey matter, we included smoothed (10mm Gaussian kernel) and unsmoothed images. We additionally included quadratic expansions of smoothed images, resulting in twelve total regressors. We regressed this model against the whole-brain hemodynamic latency map and extracted the residuals.

### Cocaine Use Disorder Dataset

For the analysis of cocaine use disorder, we analyzed a publicly available OpenNeuro Dataset ds003346^94^. The researchers collected anatomical T1w scans and 10 minutes of resting state fMRI from 74 cocaine use disorder patients (CUD) and 64 non-CUD controls as part of the Mexican MRI dataset of patients with cocaine use disorder (SUDMEX CONN). Healthy controls were matched by age (± 2y), sex, education, and handedness. Detailed study procedures can be found in prior publications using this dataset^59,60^. The MRI acquisition parameters are reported in the original study^95^.

We excluded thirty subjects recommended for exclusion in the dataset metadata due to clinical features, data collection issues, and anatomical data quality. Because these exclusions did not consider functional data quality, and motion spikes have a significant impact on resting state data^96^, we applied an additional motion exclusion criterion: Any subject with more than five frames of the resting state scan where framewise displacement exceeded 1 mm was excluded from the study, resulting in 40 CUD subjects and 39 healthy controls in our analyses (total *n* = 79). We verified that alternative motion spike thresholds produced qualitatively similar results. Data were preprocessed using the same methods as the Midnight Scan Club data described above.

For classification analyses, we tested whether the spatial pattern of latencies within the striatum predicted cocaine use disorder. We included all voxels within the Harvard Oxford striatal atlas. For each subject, we removed the mean latency across striatal voxels within each subject. This procedure ensured that the group differences in average striatal hemodynamic latency we observed did not drive the classification results. We performed leave-one-subject-out cross-validation implemented in *sklearn 1.0.2*. Following standard procedures, each voxel’s latency was mean-centered across subjects within cross-validation folds. We used logistic regression with an L1 penalty and sklearn’s default parameters. To assess the significance of our classification, we compared classification accuracy to a null distribution of classification accuracies by shuffling the group labels 1,000 times.

To analyze the spatial distributions of latencies that dissociated cocaine use disorder from healthy controls, we examined the voxel weights of the decoding model. These weights are not directly interpretable in their raw form, but they can be made interpretable by multiplying the weight vector with the data covariance matrix^97^. The resulting vector indicates a distributed pattern corresponding to the degree to which each voxel’s hemodynamic latency is associated with cocaine use disorder versus healthy control status. To test whether there were spatial gradients in this weight vector, we constructed a regression model in which the voxel weights were predicted by three regressors corresponding to x, y, and z spatial coordinates.

We compared the regression weights from this model to a null distribution of regression weights. This null distribution was constructed by shuffling the group labels in the classification models 1,000 times, extracting the voxel weights from these models, and computing the spatial regression on these voxel weights. Importantly, because only the subject group labels are shuffled, this distribution approach ensures that any spatial covariance structure that may influence the spatial pattern of voxel weights is represented in the null distribution.

## CODE AVAILABILITY STATEMENT

The data analysis code is available at https://github.com/iancballard/hemodynamic_latency.

## DATA AVAILABILITY STATEMENT

Processed data necessary for producing figures is stored alongside the analysis code at the link above. Raw data for the HCP data are available at https://db.humanconnectome.org/app. Raw data for the Midnight Scan Club (ds000224) and the cocaine use disorder dataset (ds003346) are available on https://openneuro.org/.

## ACKNOWLEDGEMENTS

We thank the D’Esposito lab for helpful discussions. Fig. 3A was created with BioRender.com.

