## Supplementary Information for "Temporal fMRI Dynamics Map Dopamine Physiology"

**Supplemental Text**

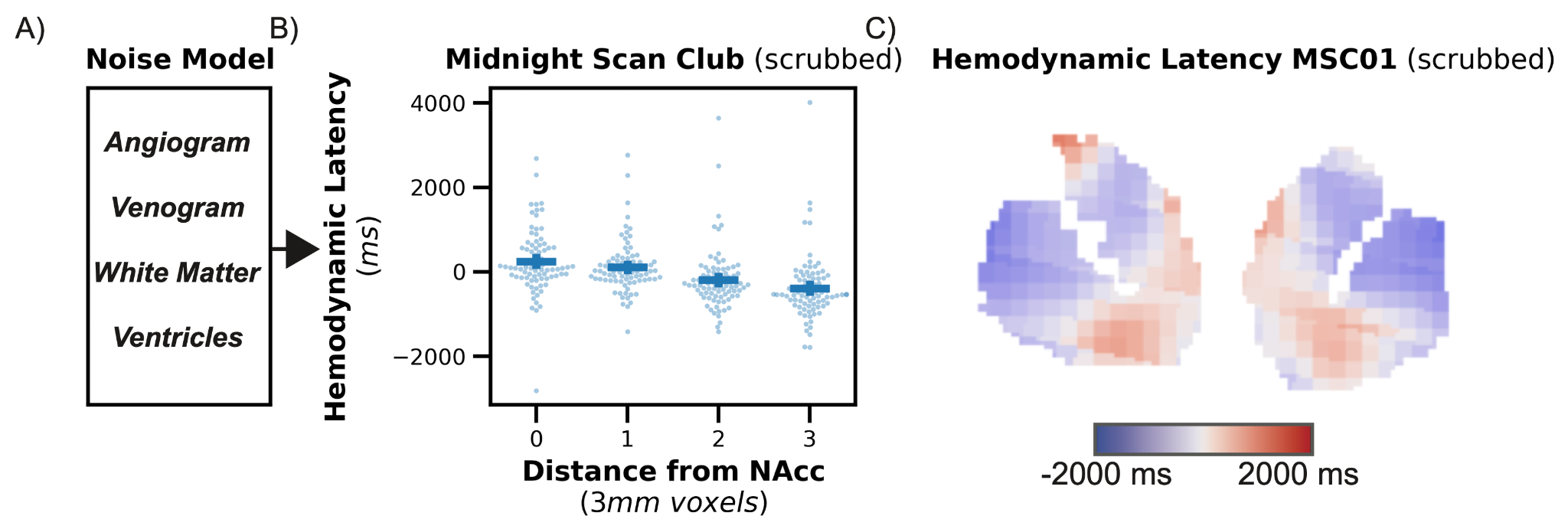

**Figure S1:** Midnight Scan Club Dataset. **A)** We constructed a noise model for individual latency data from the Midnight Scan Club dataset. **B)** Hemodynamic latencies decline at the NAcc boundary in the midnight scan club dataset in data scrubbed for sources of noise by our model. **C)** Scrubbed hemodynamic latencies for an individual MSC subject. *Error bars depict the standard error of the mean (SEM).*

*Dorsal caudate hemodynamic latency*

Across our results, we also observed increased hemodynamic latencies in the dorsal caudate (Figure 1C, D; Figure 2H). These increased lags may be due to spatial proximity to deep cerebral white matter, a region with increased hemodynamic latency^1,2^. As an exploratory analysis, we visualized the effect of spatial blurring of hemodynamic latencies from the white matter of the HCP data. This analysis accounts for spatial structure in white matter hemodynamic latencies. We observed that increased hemodynamic latencies from the deep white matter are detectable in the dorsal caudate assuming 10 mm Gaussian smoothing (Figure S1). In contrast, there was no influence on the nucleus accumbens. These results suggest that the increased hemodynamic latencies in the dorsal caudate may be driven by proximity to cerebral white matter, whereas the increased latencies observed in the nucleus accumbens are not influenced by white matter signal.

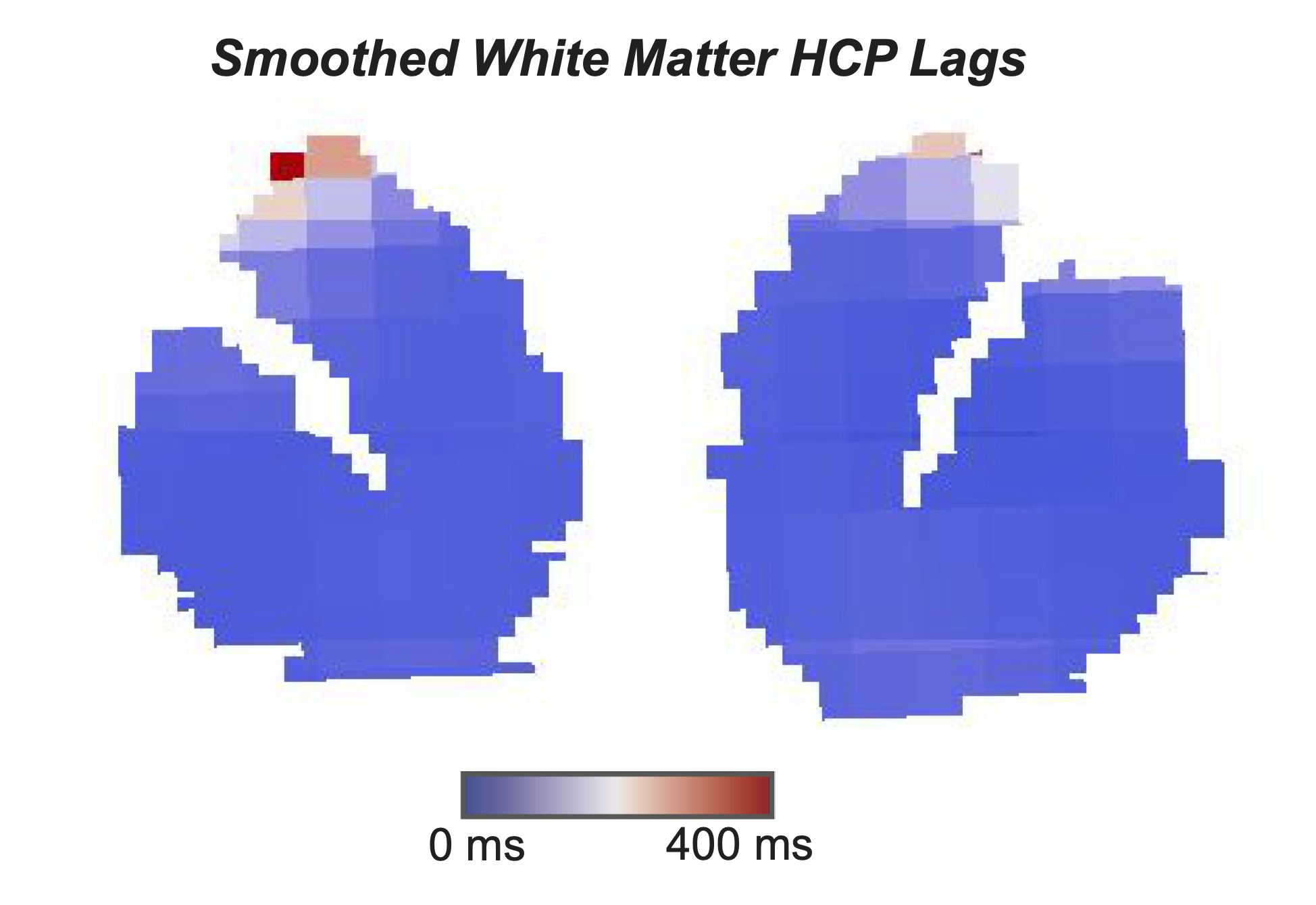

**Figure S2:** Elevated hemodynamic latencies in the white matter can influence striatal striatal hemodynamic latency estimates. Plotted is the group average hemodynamic latency plot, transformed to an example individual Midnight Scan Club subject and smoothed with a 10 mm kernel. Hemodynamic latency estimates in the dorsal caudate may be increased due to proximity to deep cerebral white matter.

*COMT relationship with hemodynamic latency*

We next assessed whether functional polymorphism in the gene encoding catechol-O-methyltransferase (COMT) was reflected in altered hemodynamic latencies. The COMT Val allele promotes increased enzymatic degradation of extracellular dopamine relative to the Met variant, resulting in lower cortical dopamine tone^3^. Because COMT is considered more critical for prefrontal than striatal dopamine clearance, we did not expect either COMT or tolcapone to influence striatal hemodynamic latencies. However, we found an interaction between tolcapone and COMT allele, *Z =* 2.2, p = .028, ∆latency= 242 ms, 95% *CI* = [26, 458] (Figure S2A), such that tolcapone reduced hemodynamic latencies more strongly for Val/Val, relative to Met/Met subjects. This effect is consistent with an U-shaped model of cortical dopamine^4^, whereby increasing cortical dopamine tone with tolcapone has opposite effects on hemodynamic latency depending on baseline dopamine tone (Figure S2D). Consistent with previous results showing the specificity of drug and gene interactions, bromocriptine, a D2 agonist, did not interact with COMT genotype, *p* > .2 (Figure S2C).

We next examined whether the COMT effects were specific to dopaminergic striatal regions. The interaction between COMT and tolcapone was not significantly different in the NAcc relative to other striatal ROIs, *ps* > .2. There was a triple interaction of region, drug, and genotype, such that the interaction between COMT and tolcapone was larger in the NAcc than S1, *Z =* -2.4, p = .017, ∆latency= -357 ms, 95% *CI* = [-652, -63]. However, there was no difference in the size of this interaction between NAcc and OTC, *p* = .139. Therefore, the effects of tolcapone and COMT genotype on hemodynamic latency in the NAcc are likely due to dopaminergic influences on systemic perfusion or downstream consequences resulting from changes in prefrontal dopamine^5^, rather than reflecting NAcc dopamine physiology.

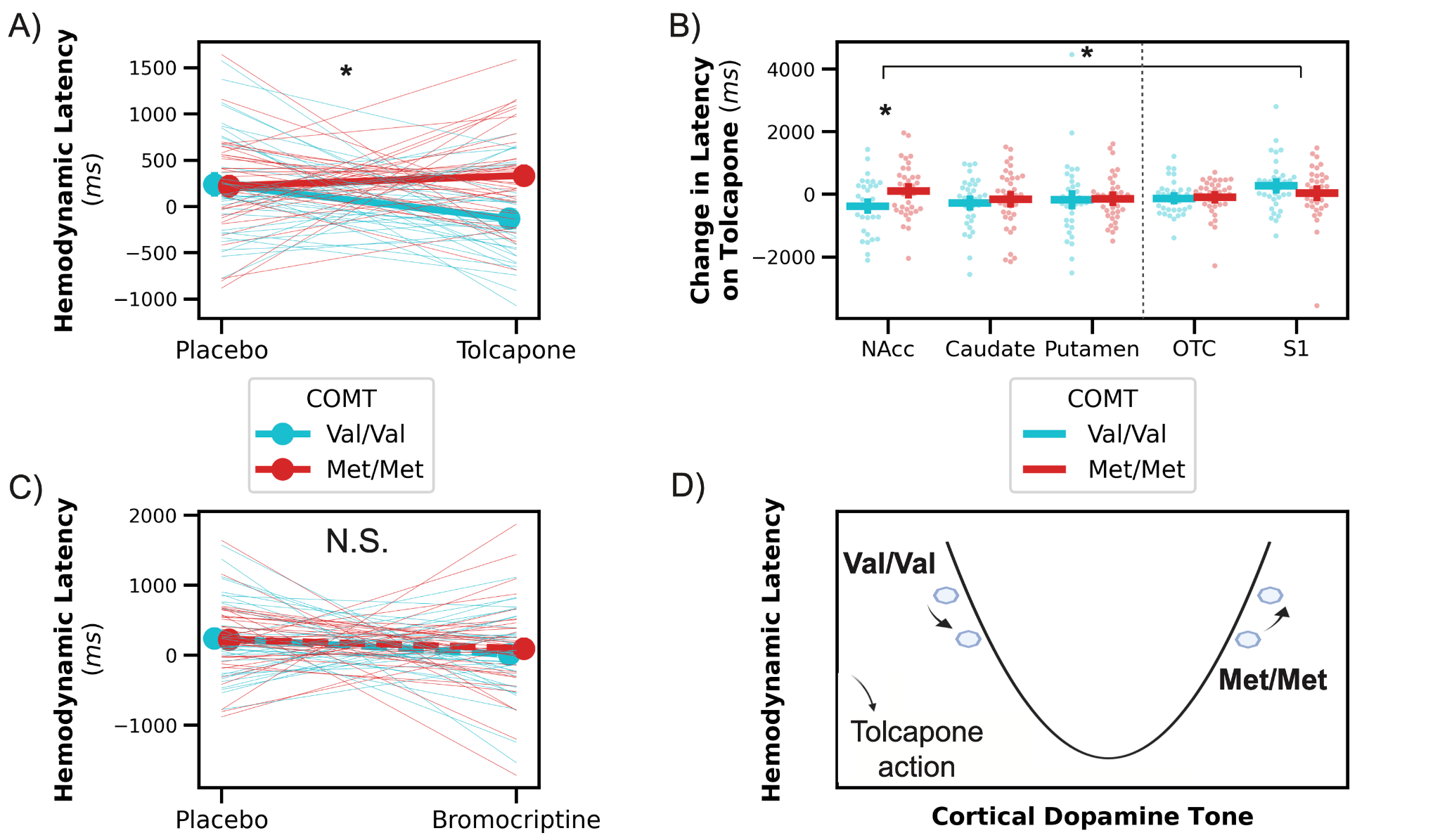

**Figure S3**: Genetic influence on hemodynamic latency. **A)** Tolcapone, a COMT inhibitor, reduces hemodynamic lags more for individuals with the COMT Val/Val genotype. **B)** The interaction between tolcapone and COMT genotype is stronger in the NAcc than in S1. However, there was no difference in this interaction effect between NAcc and OTC, indicating that this interaction may be due to extra-striatal arterial effects. **C)** Bromocriptine, a D2/3 agonist, does not interact with COMT allele to influence hemodynamic latency. **D)** The influence of tolcapone, a COMT inhibitor, on hemodynamic latency depends on COMT genotype. This reveals a U-shaped relationship between dopamine tone and hemodynamic latency. *OTC: occipitotemporal cortex; S1: somatosensory cortex*. *Error bars indicate S.E.M.*

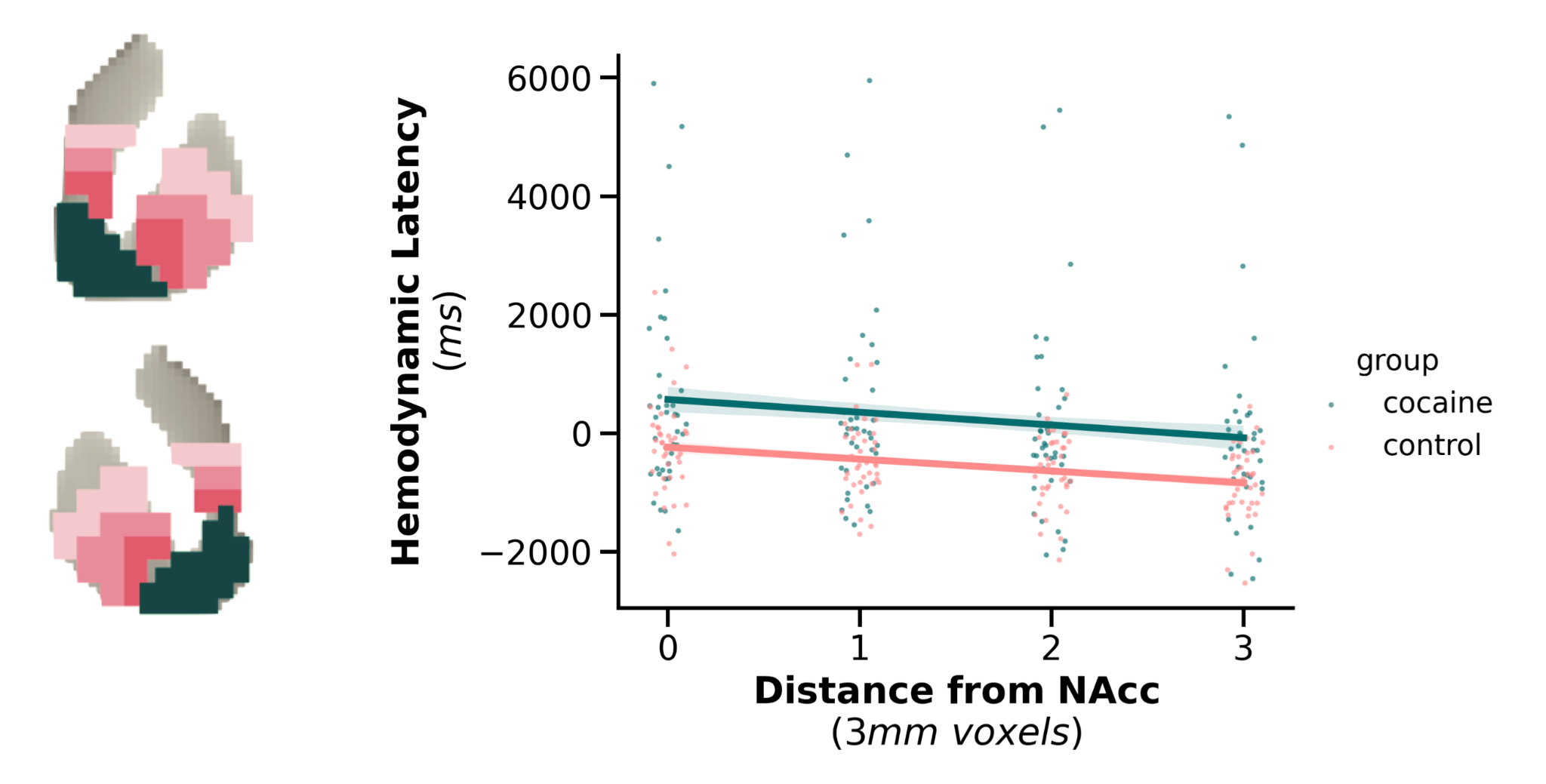

**Figure S4:** Hemodynamic latencies in the cocaine dataset trace NAcc anatomical boundaries. Hemodynamic latency in the striatum decreases linearly as distance from the NAcc increases for both the cocaine and control groups. *Error bars indicate S.E.M.*
